# An Integrated Pipeline for Cell-Type Annotation, Metabolic Profiling, and Spatial Communication Analysis in the Liver using Spatial Transcriptomics

**DOI:** 10.64898/2026.02.07.704573

**Authors:** Chi Zhang, Jun Li, Owen Luo, Xiangsheng Fu, Tallulah Andrews, Gregory R Steinberg, Dongdong Wang

## Abstract

The liver acts as a central metabolic hub, integrating systemic signals through a spatially organized pattern known as zonation, driven by the coordinated activity of diverse cell types including hepatocytes, stellate cells, Kupffer cells, endothelial cells, and immune populations. Spatial transcriptomics (ST) enables the profiling of thousands of cells with spatial resolution in a single experiment, facilitating the identification of novel gene markers, cell types, cellular states, and tissue neighborhoods across diverse tissues and organisms. By simultaneously capturing transcriptional and spatial heterogeneity, ST has become a powerful tool for understanding cellular and tissue biology. Given its advantages, there is growing demand for applying ST to uncover novel biological insights in the liver under various physiological and pathological conditions including obesity, diabetes, and metabolic dysfunction-associated steatotic liver disease (MASLD). However, to date no comprehensive and practical protocols currently exist for analyzing ST data specifically in the context of liver metabolism. Herein, we present a systematic and detailed protocol for ST data analysis using liver tissues from MASLD mouse models. This guide offers practical support for metabolic based researchers without advanced expertise in coding, mathematics and statistics enabling single-cell RNA-seq referencing for deconvolution-based annotation, curated liver cell type markers for manual annotation, and a GMT file of metabolic gene sets and flux balance analysis to analyze liver metabolic activity. This framework and integrated computational resources for decoding metabolic reprogramming and cellular heterogeneity will empower researchers to uncover novel biological pathways regulating liver metabolism in health and disease.

## Introduction

The combination of spatial organization and cell functional diversity among cells defines distinct tissues and organs that carry out specialized functions in the body. As the central metabolic hub, the liver orchestrates nutrient processing, energy homeostasis, and hormonal regulation through the coordinated activity of its diverse parenchymal and non-parenchymal cell types. Simultaneously, it serves as a key immune organ, continuously monitoring gut-derived antigens and maintaining immune tolerance while remaining capable of mounting rapid inflammatory responses. While single-cell RNA sequencing (scRNA-seq) enables the investigation of cellular functions, identities, and developmental trajectories by profiling transcriptomes at single-cell resolution^1^, it inherently loses spatial context, the organization of cells within their native environment. The spatial organization of these cells is crucial as it reveals tissue neighborhoods and local features that shape the tissue microenvironment^2^. This is especially important in the liver where the spatial arrangement of hepatocytes relative to arteries, veins, and bile ducts governs their metabolic and immune function in a pattern known as “zonation”^3^ ^4^. Importantly, this finely tuned organization is disrupted with obesity, diabetes and metabolic dysfunction-associated steatotic liver disease (MASLD), making it essential to characterize both transcriptional and spatial heterogeneity simultaneously to fully understand tissue function and disease mechanisms.

Since spatially resolved transcriptomics was crowned Method of the Year for 2020^5^ it has experienced rapid growth with a 13-fold increase in publications in the last 5 years including many involving the study of the liver^6–17^. Spatial transcriptomic technologies enable the analysis of gene expression within intact tissue architecture using methods such as microdissection-based sequencing, in situ hybridization, spatial barcoding, and probe-based RNA capture. Recent advances in these platforms have improved transcriptome coverage, sensitivity, and spatial resolution, allowing comprehensive characterization of cellular heterogeneity within complex tissues. The processing and analysis of high-throughput ST data typically involves two major stages: raw data pre-processing and downstream data analyses. Data preprocessing includes image-processing, count quantification and spatial alignment, and cell segmentation. These steps convert raw data with images into a gene-by-spot matrix with downstream analyses requiring a range of computational and statistical tools including Seurat^18–20^, Giotto^21^, or squidpy^22^. However, a major challenge for metabolism-based researchers-who often lack advanced expertise in coding, mathematics and statistics-is to apply these methods to answer their scientific questions, as currently there is not a detailed pipeline for ST analysis. For example, while pipeline have for annotating different cell types in the liver using ST have been developed^23,24^, to the best of our knowledge no integrated and systematic pipelines have been established that enable the comparison between a control and treatment group (e.g. nutritional intervention, genetic deletion, pharmacotherapy), thus limiting the practicality of using this technology for most metabolic researchers.

In this manuscript we have developed a comprehensive pipeline for ST data analysis in the context of liver metabolism and demonstrate its application using multiple datasets generated with the Visium CytAssist^©^ platform (Fig. 1). This workflow provides a detailed reference for ST data analysis comparing for metabolic-focused researchers, enabling non-technical experts in the field to harness the power of ST to uncover novel physiological regulators of metabolism and immunity as well potential novel therapeutic targets by elucidating the spatial organization of cellular and tissue biology in the liver.

**Fig. 1.**
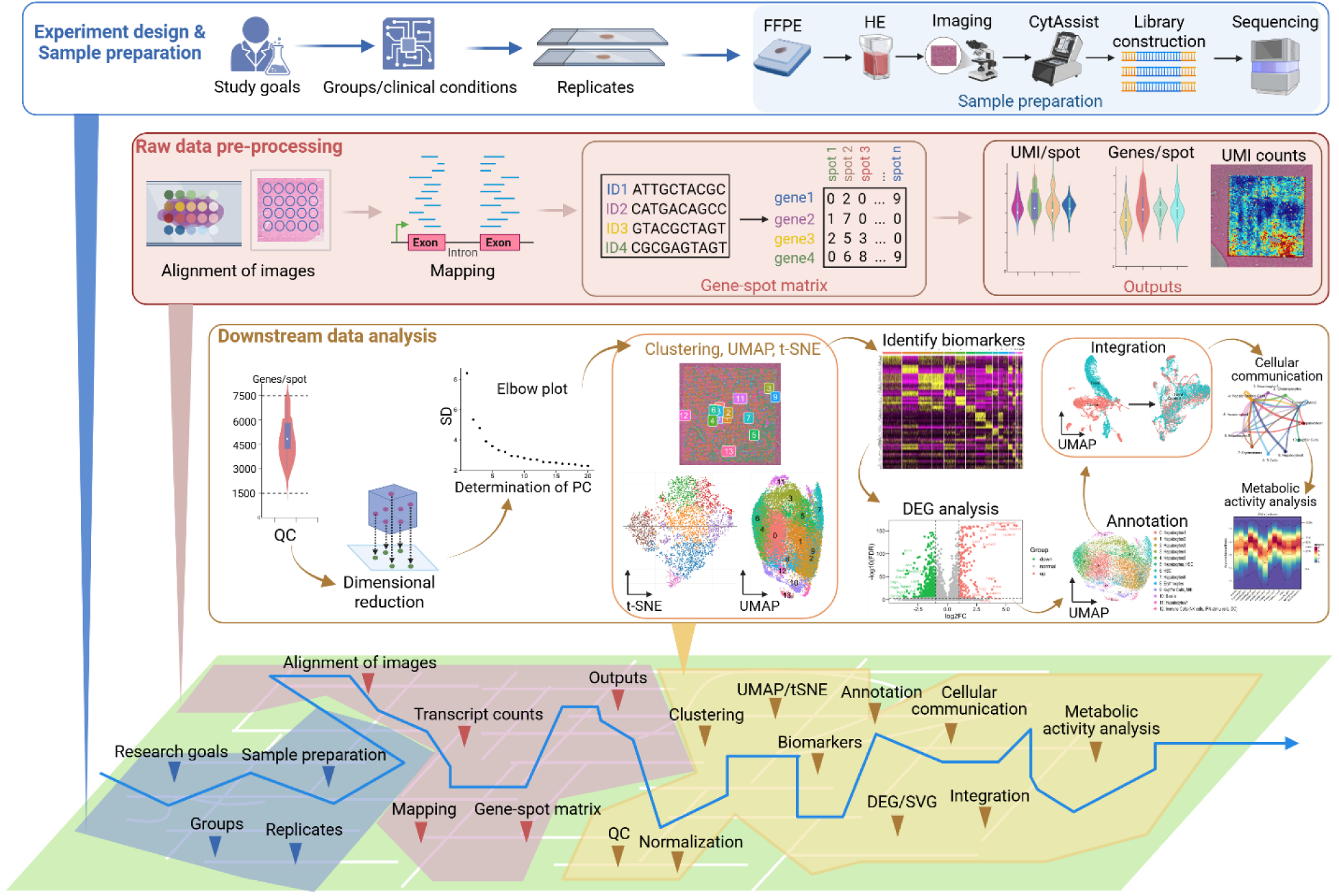
The outline of spatial transcriptomics (ST) data analysis. It includes experiment design and sample preparation, raw data pre-processing and downstream data analysis. Sample preparation comprises formalin-fixed & paraffin-embedded (FFPE) tissues preparation, H&E staining, imaging, Visium CytAssist, library construction and sequencing. Raw data pre-processing consists of alignment of images, mapping, generation of gene-spot matrix and interpretation of outputs. Downstream data analysis includes quality control (QC), normalization, linear dimensional reduction, determination of principal components (PC), clustering, non-linear dimensional reduction (UMAP/t-SNE), identification of biomarkers, differentially expressed gene (DEG) analysis, spatially variable genes (SVGs), cell type annotations, integrative analysis, pseudobulk analysis, cell-cell communication, and metabolic activity analysis.

## Results

### Experiment design and sample preparation

Experiment design is guided by specific research objectives to address key scientific questions. In this case study, we aimed to compare liver ST datasets between two conditions using a mouse model of MASLD, that closely mimics the metabolic, pathological and transcriptional characteristics of human disease^25^. Sample preparation is an important step for spatial transcriptomics, which directly influence the RNA quality. This process including paraffin-embedded formalin-fixed (PEFF) tissue preparation, HE or IF staining and imaging, as detailed in Supplementary Information.

### Data Generation and Raw data Pre-processing

The goal of data generation and raw data pre-processing is to construct a gene-by-spot expression matrix. This process includes several critical steps: sequencing, manual image alignment, read trimming, alignment of transcript reads to a reference genome, barcode correction, and probe filtering (specifically for FFPE samples) to remove off-target signals (see Supplementary Information and Fig. 1). We employed the Visium CytAssist platform, which is compatible with both formalin-fixed paraffin-embedded (FFPE) and fresh-frozen tissue samples. Unique molecular identifiers (UMIs) are counted to quantify transcript abundance, while spatial barcodes preserve transcript localization, enabling high-resolution spatial transcriptomic profiling. Raw data preprocessing is carried out using Space Ranger (a specialized pipeline developed by 10x Genomics^26^) on a computing platform that meets the minimum system requirements outlined in Box 1, as detailed in the Supplementary Information. The preparation of materials for running the Space Ranger count pipeline is outlined in Box 2. This pipeline generates a gene-by-spot expression matrix that includes transcript counts along with spatial metadata. This spatially resolved expression matrix serves as the foundation for downstream analyses. The Space Ranger count pipeline generates summary metrics for assessing sequencing quality, with interpretation guidelines provided in Box 3.

### Downstream Analysis

Downstream analysis aims to interpret spatial transcriptomic data by clustering cells, identifying cell types, and performing functional analyses, such as metabolic activity profiling, using filtered high-quality data. This process involves several key steps, including quality control (QC), normalization, linear dimensionality reduction, clustering, and visualization through Uniform Manifold Approximation and Projection (UMAP) or t-distributed Stochastic Neighbor Embedding (t-SNE). Additional steps include identifying cluster-specific biomarkers, analyzing differentially expressed genes (DEGs) and spatially variable genes (SVGs), annotating cell types, integrating data across multiple samples, conducting pseudobulk analyses, assessing cell-cell spatial communication, and evaluating metabolic activity (Fig. 2). These analyses are guided by specific research objectives and designed to address key scientific questions. In this paper, we demonstrate the application of these steps in the context of liver metabolism. Each analytical step is described in detail in the following sections.

**Fig. 2.**
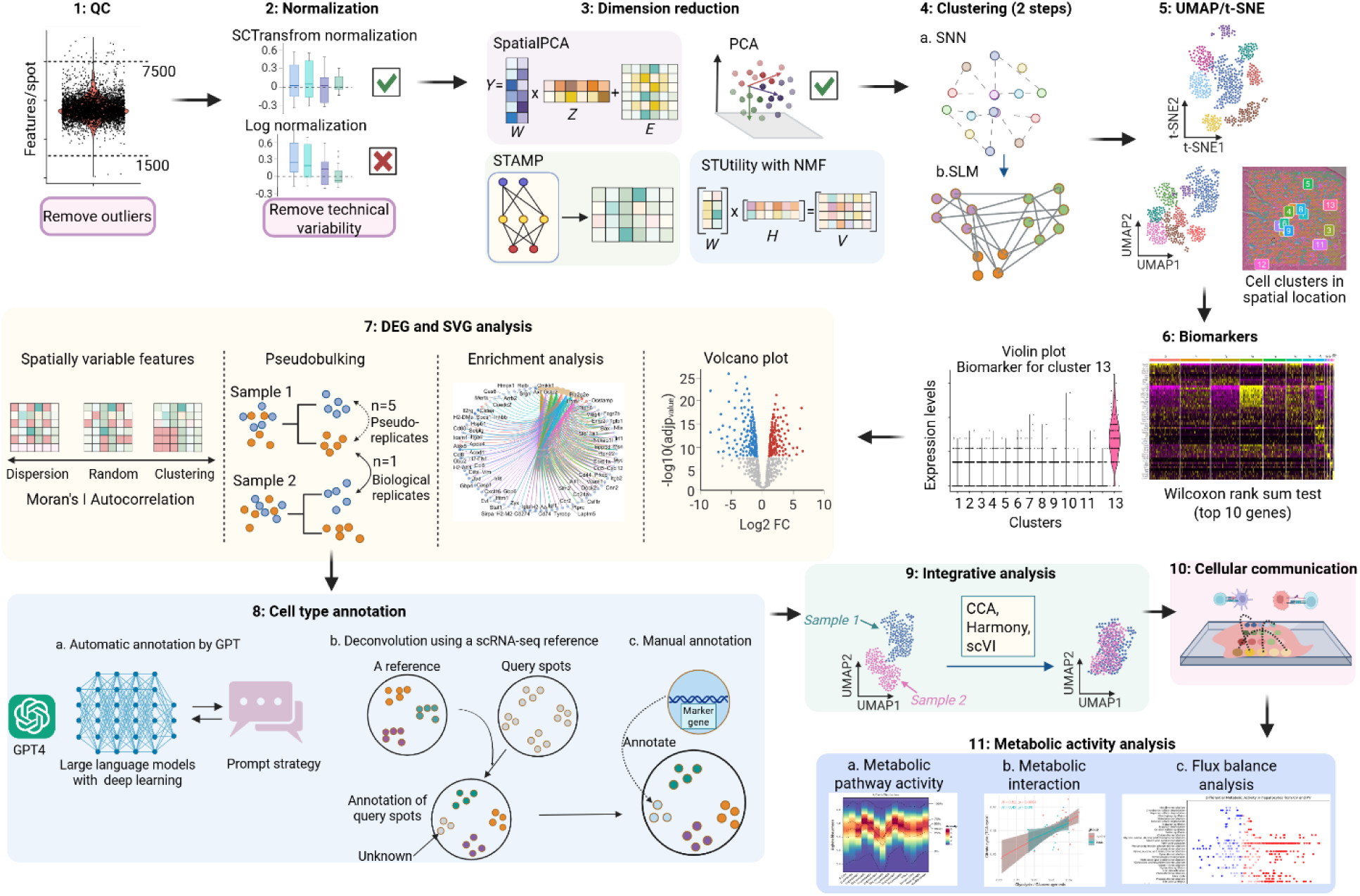
The outline of downstream analysis. This section covers quality control (QC), normalization, linear dimension reduction, clustering, non-linear dimensional reduction (t-SNE (t-distributed Stochastic Neighbor Embedding) and UMAP (Uniform Manifold Approximation and Projection)), identification of cluster biomarker, cell type annotation (both automated and manual), differentially expressed gene (DEG) analysis, spatially variable genes (SVGs) analysis, integrative analysis across multiple samples, cellular communication and metabolic activity analysis.

### Quality Control (QC) and normalization

Quality control (QC) ensures the reliability and accuracy of ST data by identifying and removing low-quality observations and outliers. Spots with very low gene counts typically reflect low-quality regions or partial tissue coverage. QC is performed by filtering spots based on the number of unique genes detected. Based on the distribution (Fig. 3A), spots with fewer than 1,500 and more than 7,500 unique features are filtered out, as detailed in Supplementary Information. It is important to note that current QC approaches, based on fixed or data-driven global thresholds, often overlook biological heterogeneity across different tissue regions, which may introduce bias. As such, if manual inspection of tissue segmentation confirms accurate spot selection, it may be appropriate to forgo QC filtering altogether in certain ST datasets^27^.

**Fig. 3.**
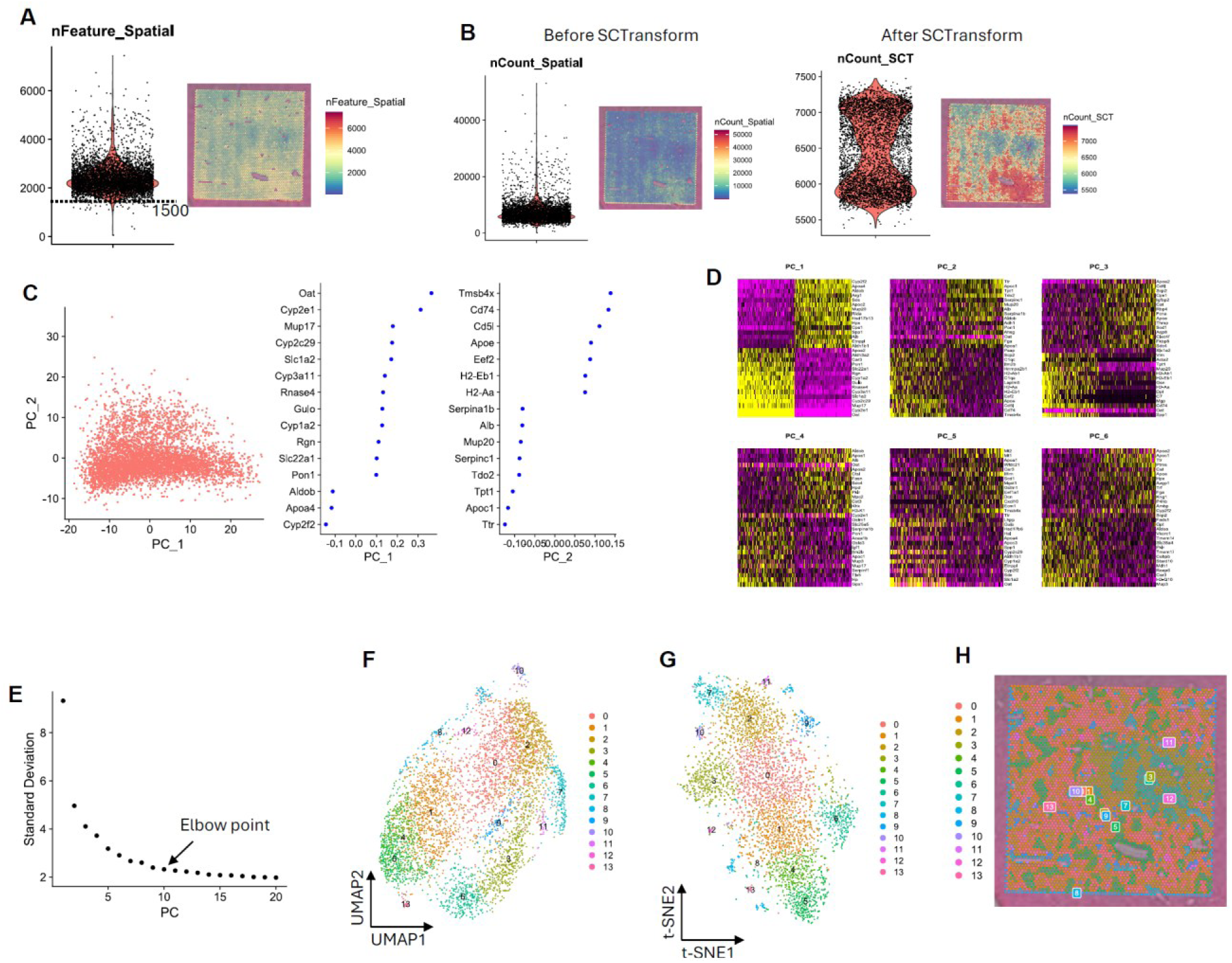
Quality control (A), normalization (B), dimension reduction using PCA (C), DimHeatmap (D) and elbow plot (E) to determine which PCs are used for following analyses, visualization of clusters in UMAP and t-SNE (F, G), and spatial localization of clusters on the tissue slide (H).

Following quality control, normalization is applied to correct for technical variability, such as differences in sequencing depth and molecule counts per spot, while preserving true biological differences, including variations in RNA content and cell density. Sctransform is a robust normalization method well-suited for handling tissue samples with variable mRNA content driven by technical factors^28^. In this protocol, we use sctransform to normalize the data, identify highly variable features, and scale the dataset for downstream analysis, as detailed in Supplementary Information. This step returns datasets including normalized and scaled data suitable for following dimension reduction (Fig. 3B).

### Dimension reduction

To extract biologically meaningful signals and reduce noise, dimensionality reduction is commonly applied to transform high-dimensional data into a low-dimensional representation^29,30^. This step is essential for effective visualization, interpretation, and a range of downstream analyses. PCA is applied on the normalized and scaled data to do dimensional reduction, as detailed in Supplementary Information. The variable features identified by sctransform are used as input for dimensionality reduction. The output includes a list of genes with the most positive loadings for each principal component, representing gene modules that exhibit correlated expression patterns across spots in the dataset (Fig. 3C), and heatmap was used to determine which PCs capture the major sources of variation (Fig. 3D). An Elbow Plot was used to select the optimal number of PCs for clustering. In our datasets, the elbow is observed around PC9-10, suggesting that the majority of biologically meaningful signals are captured within the first 10 PCs (Fig. 3E). Data after dimension reduction can be used for following clustering.

### Clustering

Clustering is a critical step in the ST data analysis workflow, aimed at grouping spots based on similarities in gene expression profile and spatial context (Fig. 2). Clustering is performed in two main steps: 1) construct a SNN graph, and 2) iteratively grouping of cells into clusters, as detailed in Supplementary Information. The group of spots can be visualized using UMAP or t-SNE.

### Non-linear dimensional reduction (UMAP/t-SNE)

Visualization of the identified clusters in a two-dimensional space is important to interpret results. This step was performed using non-linear dimensionality reduction techniques (Fig. 2). These methods project high-dimensional data onto lower-dimensional manifolds while preserving the intrinsic structure of the data. Two visualization methods t-SNE (t-distributed Stochastic Neighbor Embedding)^31^ and UMAP (Uniform Manifold Approximation and Projection)^32^ were used in this workflow. UMAP constructs a topological representation of the high-dimensional data and projects it in a way that better preserves both local and global relationships, compared to t-SNE. In this protocol, UMAP and t-SNE are displayed as two-dimensional scatter plots, where each point represents a spatial capture spot (Fig. 3F and G). The positioning of each spot is based on its low-dimensional embedding generated by UMAP or t-SNE, and points are color-coded according to their assigned clusters, as detailed in Supplementary Information. In addition, clusters can be visualized in their anatomical context, which overlays cluster identities on the original tissue image (Fig. 3H).

### Identification of cluster biomarkers

Identification of cluster-specific biomarkers is essential for accurate cluster annotation and helps refine the focus of subsequent research. Molecular biomarkers also provide insights into the current functional states of specific cells or tissues. ST enables the discovery of biomarkers within complex tissue environments by preserving spatial context. In ST, biomarker analysis can reveal both anatomical and microanatomical features by identifying markers that are specific to defined spatial clusters. These biomarkers are typically detected through DEG analysis, which identifies genes that are significantly upregulated in one cluster relative to all others (Fig. 2), as detailed in Supplementary Information. The top 10 markers for each cluster are visualized in a heatmap (Fig. 4A). Expression patterns of selected marker genes, such as those for clusters 4 and 5, are displayed using violin plots (Fig. 4B). These markers can also be spatially visualized on the UMAP (Fig. 4C), or directly on the tissue image (Fig. 4D). The identified cluster biomarkers are used for subsequent cell type annotation.

**Fig. 4.**
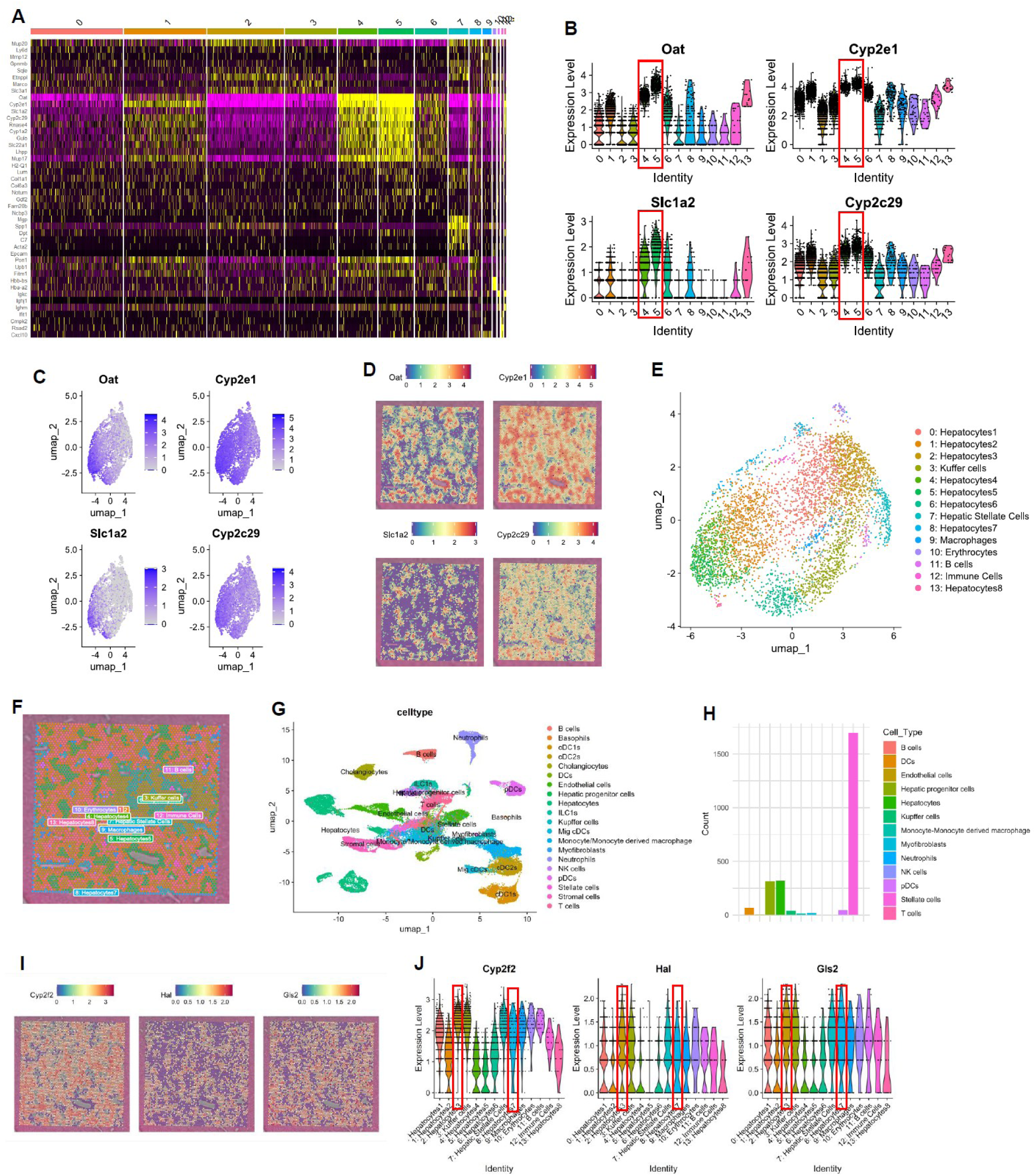
Determination of cluster biomarkers and cell type annotations. A, Heatmap showing the top 10 biomarkers for each cluster. B, Violin plot displaying the expression of marker genes for clusters 4 and 5. C & D, Visualization of marker gene expression for clusters 4 and 5 on UMAP (C) and overlaid on the H&E-stained tissue image (D). E & F, Automated cell type annotation results from GPT-4 shown on UMAP (E) and in the H&E image (F). G. Cell type annotation based on the integrated scRNA-seq reference, visualized on UMAP. H. Cell type annotation after deconvolution using the generated scRNA-seq reference, visualized on UMAP. I. Spatial visualization of periportal hepatocyte marker expression overlaid on the H&E-stained tissue image. J. Violin plot showing the expression of periportal hepatocyte markers.

### Cell type annotation

Cell type annotation is a fundamental but very challenging step in ST data analysis^33^. Accurate annotation reveals the spatial organization of cell types, characterizes tissue-specific biological features, and enhances our understanding of cellular and tissue functions in both health and disease^34^. In this case study, we apply a three-step workflow for cell type annotation: automatic annotation using GPT4^33^, deconvolution with a scRNA-seq reference, and manual cell type annotation^35^.

#### a. Automatic annotation using GPT

Annotation is a labor intensive and time-consuming process. To streamline this step, we apply automatic cell type annotation as an initial approach. In this protocol, we use GPTCelltype^33^, an R package that leverages GPT, a class of large language models (LLMs), to perform automated annotation. GPT is a prominent framework for generative artificial intelligence built on deep neural networks. GPT-4 in particular has demonstrated the potential to annotate cell types across a wide range of tissues due to its extensive training on diverse datasets^33^. We utilize GPTCelltype to automate cell type annotation by inputting identified cluster biomarkers along with a basic prompt strategy^33^ (Fig. 2), as detailed in Supplementary Information. The annotation results are visualized on the UMAP (Fig. 4E) and can also be mapped onto the original tissue image (Fig. 4F). In this automatic annotation, small cell populations, such as hepatocyte subtypes, dendritic cells (DCs), neutrophils, and natural killer (NK) cells, could not be reliably identified. Therefore, to refine and validate these annotations, we apply deconvolution using a scRNA-seq reference in the following sections. Although we use GPT-based annotation primarily as an initial step in our protocol, such approaches are poised to shift annotation workflows from manual or semi-automated processes toward fully automated and scalable solutions^33^.

#### b. Annotation via deconvolution using a scRNA-seq reference

The transcriptional profile of each spot represents a mixture of transcripts from multiple cells and cannot capture the true expression profile of individual cells. To address this limitation, deconvolution is used to estimate the cell type composition within each spot. To infer the cellular composition of each spot, cell type deconvolution was employed to predict the underlying mix of cell types^36,37^. A high-quality, tissue-matched scRNA-seq reference is critical for accurate deconvolution in ST analysis (Fig. 2). We generated a high-quality scRNA-seq reference specific to mouse livers with MASLD^38–41^ (see Supplementary Information) as visualized on a UMAP plot (Fig. 4G), which is critical for ensuring accurate deconvolution. In this protocol, we applied the SpatialDWLS method within the Giotto framework as well as RCTD^42^, using scRNA-seq data as a reference to improve cell type annotation, as detailed in Supplementary Information. RCTD improved the identification of small cell populations; however, it partially misclassified stellate cells as hepatocytes (Fig. 4H). To validate annotation using deconvolution, we further performed manual annotation.

#### c. Manual annotation

Automatic annotation and deconvolution often yield conflicting or incomplete cell labels, therefore, manual annotation is usually required to validate these initial annotations before proceeding with downstream analyses^43,44^. In this protocol, we perform manual annotation at the cluster level using marker-based strategies, relying on cluster-specific biomarkers identified in earlier steps (Fig. 2). For manual annotation, we generated a curated reference of liver cell type markers according to the literatures^38–41^ and databases^45–47^, as detailed in Supplementary Information. Expression of biomarkers (*Oat, Cyp2e1, Slc1a2*) for peri-central hepatocytes were spatially visualized on the tissue and violin plot (Fig. 4B and 4D). The periportal hepatocyte markers *Cyp2f2, Hal* and *Gls2* were visualized as shown in Fig. 4I and 4J.

Cell type annotation poses several challenges, including the identification of novel cell types and clusters expressing markers from multiple cell types. Therefore, it is essential to validate annotation labels using multiple independent approaches and, when possible, consult domain experts.

### DEG analysis, pathway enrichment analysis and Spatially Variable Genes (SVGs) analysis

To evaluate gene expression changes either between clusters or within the same clusters under different conditions, highly variable genes (HVGs)^48^ (also named as DEG) analysis between cell types is performed, as detailed in Supplementary Information. In this case study, we compare hepatocyte 1 and hepatocyte 2 as an example. The resulting DEGs are visualized using a volcano plot (Fig. 5A). To investigate functional differences, GO enrichment analysis is conducted to identify significantly enriched pathways (Fig. 5B).

**Fig. 5.**
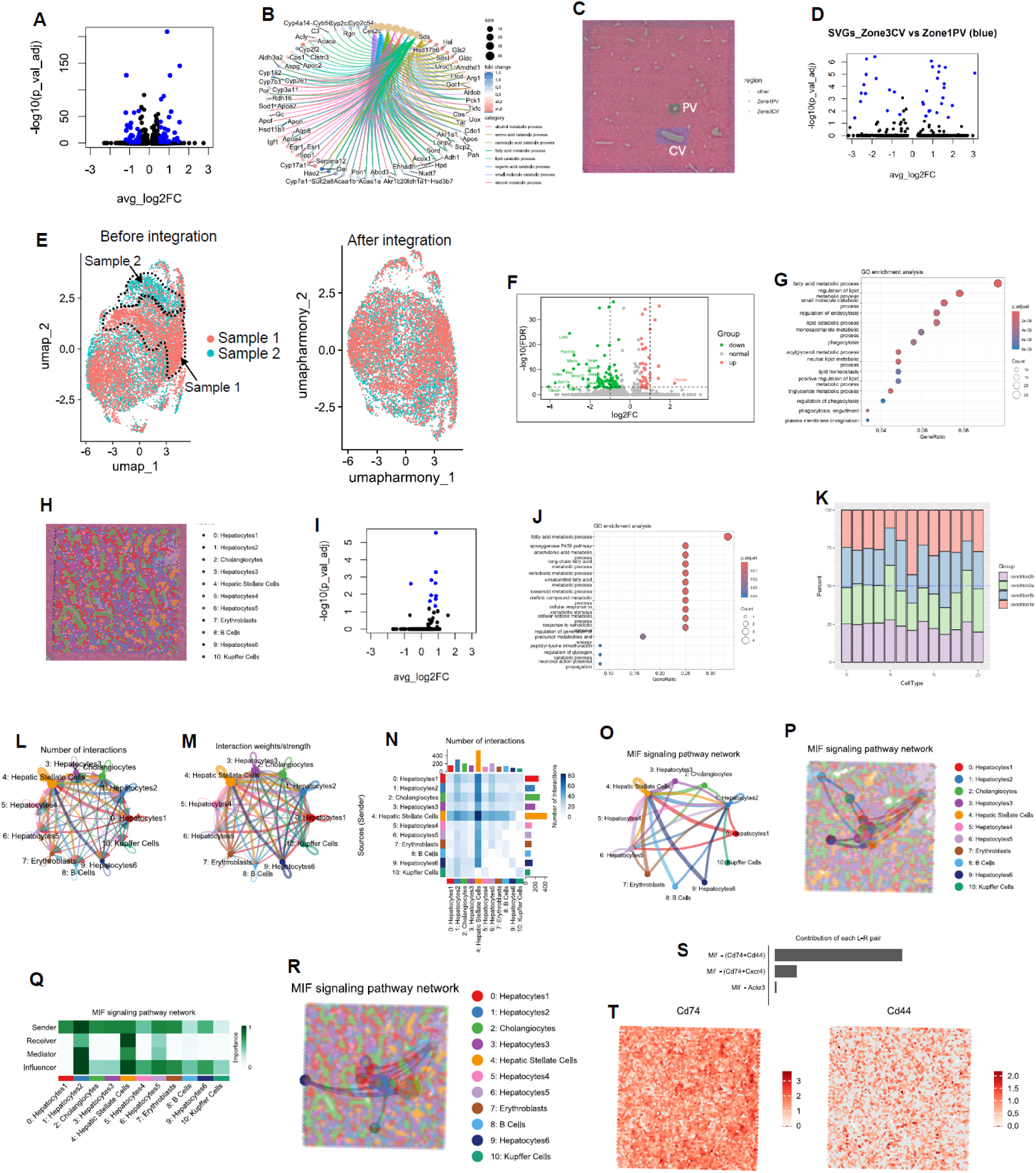
DEG analyses and cell-cell spatial communication. A. DEG analysis between clusters. B. GO enrichment analysis based on cluster-specific DEGs. C. Extraction of regions of interest surrounding the central vein (CV) and portal vein (PV). D. DEG analysis comparing cells located near the CV and PV. E. Integrative analysis across multiple samples. F–G. Following integration, identification of DEGs across conditions (F), GO enrichment analysis between conditions (G). H. UMAP visualization of samples after integration, annotation, and dataset splitting. I–J. Following pseudobulking, identification of DEGs after pseudobulking (I), GO enrichment analysis between conditions (J). K. Bar plot showing cell type proportions in tissue. L-M. Circle plots showing the number (L) and strength (M) of interactions between different cell groups. N. Heatmap displaying the intensity of cell–cell interactions. O-P. Visualization of macrophage migration inhibitory factor (MIF) signaling using a circle plot (O) and spatial tissue image (P). Q-R. Heatmap (Q) and spatial image (R) showing the importance of the MIF signaling pathway across cell types based on network centrality scores. S. Contribution of individual ligand–receptor pairs to the MIF signaling pathway. T. Spatial expression and localization of representative ligand (Cd44) and receptor (Cd74) in liver tissue sections.

Advances in ST have made it possible to identify genes whose expression patterns exhibit spatial structure. These spatial patterns may reflect communication between adjacent spots, local microenvironmental influences, or migration of specific cells populations to defined anatomical regions^49^. Different from highly variable genes (HVGs) without incorporating spatial information, spatially variable genes (SVGs) analysis is capable to capture spatially patterned expression, while also helping reduce the dimensionality of spatial transcriptomics data^50^. In this case study, we describe two complementary workflows to identify molecular features associated with spatial location. The first approach involves performing DEG analysis based on pre-annotated anatomical regions, guided by prior knowledge of tissue structure. For example, we selected regions of interest surrounding the central vein (CV) and portal vein (PV) (Fig. 5C) and performed DEG analysis between these regions (Fig. 5D), as detailed in Supplementary Information. An alternative approach is annotation-free, which detects spatial expression patterns directly from the data. In this case, we were unable to detect SVGs that strongly correspond to the microscopic anatomical structure of the liver (Extended Data Fig. 1E), unlike what is commonly observed in the brain^51^.

### Integrative analysis across multiple samples or conditions

Integrative analysis aims to remove batch or condition-specific effects while preserving true biological variation, the evaluation of cell-type-specific responses to different stimuli^52,53^. In this case study, without integration, clustering results are confounded by both cell type and experimental conditions (Fig. 5E), making downstream comparisons difficult. We perform integration using sctransform-normalized data and apply both Harmony and CCA to harmonize datasets across conditions, as detailed in Supplementary Information. Prior to integration, the two samples appear as distinct clusters; after integration, the samples are well-aligned and overlap substantially, indicating effective correction of batch or condition-specific effects and successful integration of shared biological signals (Fig. 5E). We further performed DEG analysis (Fig. 5F) and GO enrichment analysis (Fig. 5G) on hepatic stellate cells (HSCs) between two different conditions following integrative analysis. These analyses can be extended to any cell type of interest.

### Pseudobulk analysis

The DEG analysis without pseudobulking often results in many false positives^54^, as it treats each spot within the same sample as an independent biological replicate. To avoid false conclusions in DEG analysis, we performed differential expression analysis using a pseudobulk approach, treating samples, not individual spots, as biological replicates, as detailed in Supplementary Information. This reduces false positives and enables more reliable statistical inference. The pseudobulk approach requires a minimum of three biological replicates per condition, although more replicates are strongly recommended to increase statistical power. In this case study, the resulting DEGs after pseudobulking are visualized using a volcano plot (Fig. 5H). Functional differences between conditions are identified using GO enrichment analysis, with the results presented in a dot plot (Fig. 5I).

### Quantification of cell type composition

To assess the relative abundance of different cell types within the tissue, we count the number of cells per cell type by extracting this information from the meta data. The proportions of each cell type are displayed in a bar plot (Fig. 5J). It is important to interpret these counts with caution, as the observed cell type proportions may be influenced by the specific tissue regions selected for analysis.

### Cellular communication

Cell-cell communication is critical to coordinate diverse biological processes within tissue niches, such as immune responses, cellular metabolism and organ development and is central to understanding the complexity of the liver microenvironment^55,56^. Specifically, a key advantage of ST is its ability to dissect with spatial resolution, this cellular communication involving not only paracrine signaling but also autocrine mechanisms, tight junctions and mechanical effects^57^. In this case study, we apply CellChat, a tool that leverages network analysis and pattern recognition techniques, to investigate cellular communication in liver tissues, as detailed in Supplementary Information. First, all samples were integrated through the standard integration workflow, followed by cell type annotation and splitting into individual datasets for analysis (Fig. 5K). The number and strength of interactions between cell types are visualized using circle plots (Fig. 5L, 5M) and heatmaps (Fig. 5N). Cell-cell communication networks are organized by signaling pathway. For example, we highlight the macrophage migration inhibitory factor (MIF) signaling pathway by visualizing its network using a circle plot and a spatial image overlay (Fig. 5O, 5P). To evaluate the influence of each signaling pathway across cell types, network centrality scores are computed and displayed in a heatmap (Fig. 5Q) and spatially visualized in tissue images (Fig. 5R). To identify key mediators within a signaling pathway, the contributions of individual ligand-receptor pairs are quantified (Fig. 5S). The expression levels and spatial distributions of representative ligands (e.g., *Cd44*) and receptors (e.g., *Cd74*) are visualized directly in tissue sections (Fig. 5T). These spatially resolved cell-cell communication analysis provides a powerful framework for exploring the complexity of these interactions *in situ* not possible using other methodologies.

### Metabolic activity analysis

#### a. Metabolic pathway activity

Cellular metabolism is central to cell functions, and the liver is the body’s central metabolic hub^58^. The metabolism of carbohydrates, lipids, and amino acids underpins energy production, biosynthesis, and signaling processes across all physiological systems^58^. Metabolic activity can be evaluated using Gene Set Enrichment Analysis (GSEA)^59^ with predefined gene sets associated with specific metabolic pathways. Refined and comprehensive metabolic gene sets for mouse models remain limited. In this protocol, we firstly developed mouse-specific metabolic gene sets derived from human KEGG pathways and other metabolic pathways in the liver (see Supplementary Information). To quantify pathway activity, we applied integrated gene set enrichment analysis (irGSEA)^60^, as detailed in Supplementary Information. This analysis can reveal differences in metabolic activity patterns across liver cell types, as illustrated in the heatmap (Fig. 6A) and dot plot (Fig. 6B). To explore pathway-gene relationships, we generated a cnetplot that visualizes the network structure linking genes to their associated metabolic pathways (Fig. 6C) based on GESA. To measure the activity of specific pathways, such as fatty acid oxidation and the TCA cycle, we generated density heatmaps (Fig. 6D) and violin plots (Fig. 6E). Researchers can assess the activity of specific pathways of interest simply by modifying the pathway names.

**Fig. 6.**
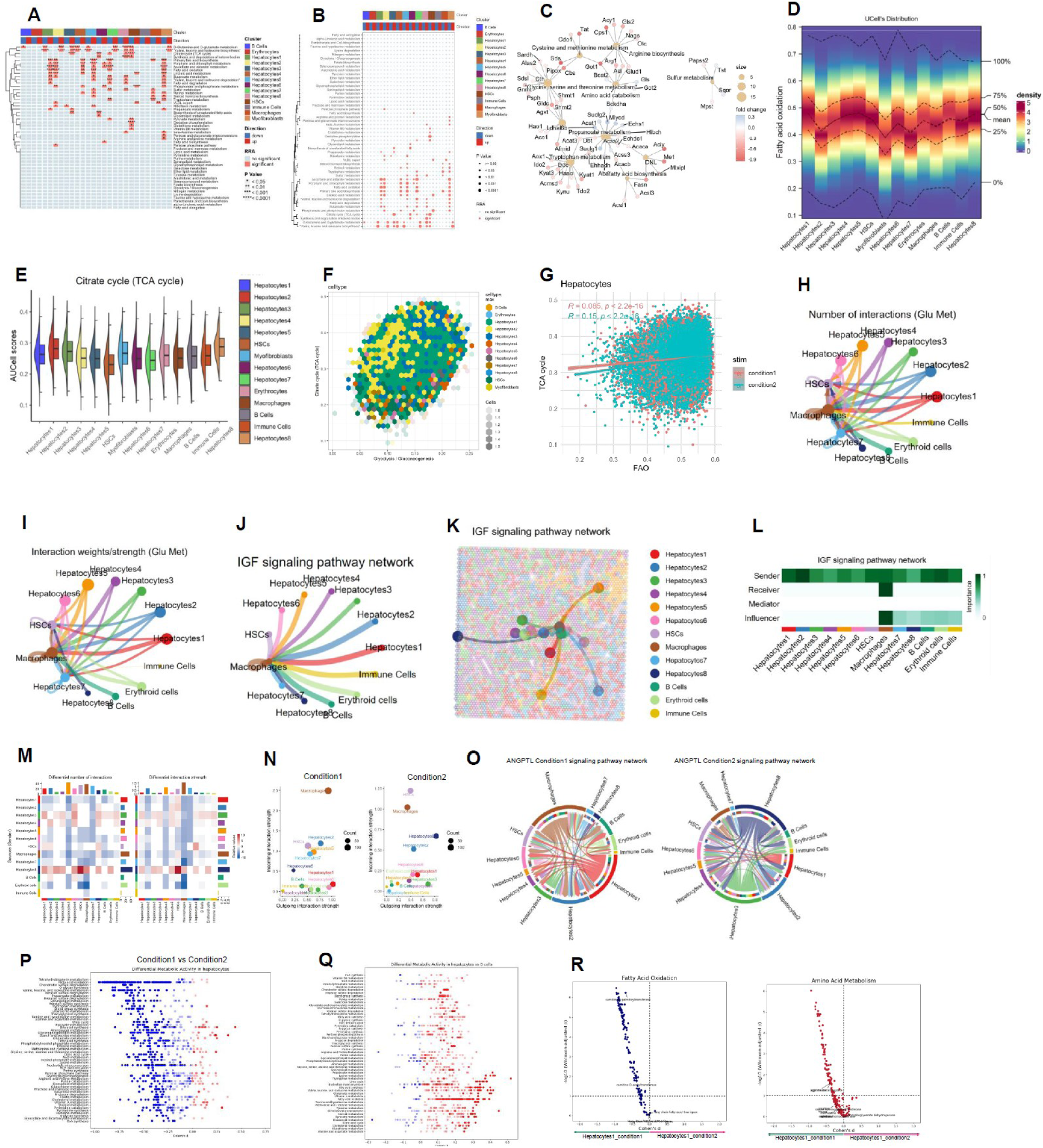
Metabolic Activity Analysis in the Liver. A–B. Heatmap (A) and dot plot (B) showing differences in metabolic activity patterns across liver cell types using irGSEA. C. Cnetplot illustrating core genes associated with the top regulated metabolic pathways. D. Density heatmaps depicting specific metabolic activities across cell types. E. Violin plot displaying metabolic activity of selected pathways across cell types. F. Correlation heatmap showing metabolic interactions between different pathways across cell types. G. Differences in metabolic pathway interactions within specific cell types. H, Number of interactions related to glucose metabolism. I, Total interaction strength (weights) in glucose metabolism. J, IGF signaling pathway networks. K, Spatial visualization of metabolic communication based on the IGF signaling pathway. L, Network centrality scores for the IGF signaling pathway. M. Heatmap showing differences in the number and strength of glucose metabolism-related interactions between two conditions. N. Scatter plot identifying cell populations with significant changes in signal sending or receiving. O. Chord diagram illustrating alterations in signaling networks between conditions. P. Comparison of metabolic program activity between two conditions using flux balance analysis (FBA). Q. FBA-based comparison of metabolic program activity across different cell types. R. Volcano plots showing differential metabolite activity in fatty acid oxidation and amino acid metabolism between conditions.

#### b. Metabolic interactions

Metabolic interactions are critical for coordinating cellular function, because cells exchange nutrients, compete for metabolic substrates, and secrete metabolites that influence the function of neighboring cells. These interactions reshape cellular behavior, modulate the tissue microenvironment, and define immune niches. In this case study, we introduced two approaches to assess metabolic interactions, including correlation analysis of metabolic pathway activity scores from ssGSEA and intercellular communication via ligand-receptor gene pairs associated with metabolic processes via CellChat. For example, we measured the correlations between glycolysis/gluconeogenesis and TCA cycle in hepatocytes (Fig. 6F) and also compared pathway correlations under different conditions (Fig. 6G), as detailed in Supplementary Information. Furthermore, we also curated ligand-receptor interactions associated with glucose, lipid, and amino acid metabolism from the CellChat database to perform cell-cell communication analysis using CellChat. We quantified the number of interactions (Fig. 6H) and total interaction strength (Fig. 6I) in glucose metabolism-related signaling networks. Specific signaling pathways, such as IGF signaling, were further analyzed using dot plots (Fig. 6J) and spatial mapping (Fig. 6K). We also can perform network centrality analysis to identify the most influential cell types and pathways (Fig. 6L). Researchers can easily assess interactions between metabolic pathways by simply modifying the metabolic names. To investigate whether metabolic communication differs between conditions (e.g., control vs. treatment or healthy vs. diseased), we demonstrate how to compare intercellular interactions using lipid metabolism as an example. We assessed both the number and strength of interactions to determine the overall impact of stimulation (Fig. 6M). Metrics were analyzed across cell types to identify which populations were most affected (Fig. 6N). Chord diagrams were used to visualize alterations in communication networks, highlighting multiple ligand-receptor pairs contributing to differences in intercellular communication between different cell types (Fig. 6O).

#### c. Flux balance analysis (FBA)

Given the complexity and connectivity of metabolic networks^61^, FBA has become a widely used computational method for investigating global metabolic circuits. FBA predicts the flow of metabolites through a biological system using a genome-scale metabolic model (GEM). In this case study, we performed FBA using Compass^62^ with RECON2 as a GEM, as detailed in Supplementary Information. Downstream analysis of the Compass results was then conducted to identify condition- and cell-type-specific metabolic changes, as detailed in Supplementary Information. Differential activity of individual metabolic reactions between conditions (Fig. 6P) and among cell types (Fig. 6Q) was assessed. Additionally, differences in pathway-level metabolite activity, such as in fatty acid oxidation and amino acid metabolism, were visualized using volcano plots (Fig. 6R). This approach enables robust inference of metabolic activity in the mouse liver using spatial transcriptomics data.

## Concluding Remarks

In this protocol, we present a comprehensive and practical pipeline for ST analysis tailored specifically to the study of liver metabolism in preclinical models of MASLD. By integrating data preprocessing, multi-tiered cell type annotation, differential gene expression, spatially resolved metabolic pathway analysis, and advanced cell-cell communication modeling, our framework empowers metabolic researchers to extract biologically meaningful insights from ST data. The use of three complementary annotation strategies, including GPT-based automated, deconvolution-based, and manual, ensures robust and accurate cell-type classification, while the integration of metabolic gene set enrichment and flux balance analysis (FBA) offers novel approaches for dissecting the spatial metabolic landscape at high resolution.

Our pipeline enables a unique interrogation of spatial metabolic heterogeneity, zonal dysregulation, and metabolic-immune cross-talk that underlie MASLD pathogenesis. The capacity to resolve metabolic pathway activities, quantify metabolic interactions, and infer cell-cell communication within the spatial context of intact tissue architecture represents a significant advance over conventional transcriptomic or single-cell approaches. Furthermore, this workflow is accessible to metabolic investigators who may not possess extensive computational expertise, thus democratizing the use of ST in liver metabolism research. Importantly, while the protocol described here uses mouse liver from a MASLD model, its flexible and modular design makes it adaptable for a wide range of research applications, including other metabolic diseases, pharmacological intervention studies, genetic models, and potentially also human clinical samples.

## Limitations and Future Directions

While the proposed pipeline provides a comprehensive framework for ST analysis in liver metabolism, several limitations should be acknowledged. First, the current analysis is based exclusively on a mouse model of MASLD, and validation in human liver tissues will be critical to assess translational relevance. Second, the resolution of current ST platforms, such as Visium CytAssist, remains limited to capturing spot-level transcriptomes rather than true single-cell resolution, which may obscure the precise characterization of rare or transitional cell states. Third, although multiple cell type annotation strategies were employed, including GPT-based, deconvolution, and manual approaches, certain small cell populations remained challenging to resolve, emphasizing the ongoing need for improved automated annotation methods. Fourth, while bioinformatic analyses identified numerous differential gene expression patterns and signaling interactions, experimental validation through orthogonal approaches such as immunohistochemistry or spatial proteomics remains essential to confirm key findings. Finally, larger sample sizes with increased biological replicates will be important to enhance statistical power for detecting subtle but biologically relevant differences, particularly in pseudobulk analyses. Moving forward, integration with emerging high-resolution spatial platforms, multi-omics modalities, and longitudinal studies will further advance our understanding of the spatial metabolic microenvironment and its role in regulating liver metabolism, disease progression and therapy responses.

## Data availability

The example files including FASTQ files, brightfield images, JSON files, HDF5 format files, a spatial folder (spatial/), a scRNA-seq reference for cell type deconvolution, RDS file for pseudobulk analysis for this protocol are available at: https://www.dropbox.com/scl/fo/occj7x04tbkjrnqfrko1b/AOV196ag11OrAIUtWagY6g8?rlkey=qyzhi6f20x517znqovu4n8q22&st=j4mhkfpm&dl=0. The summary metrics (web_summary.html file), a marker reference for manual annotation, refined and comprehensive metabolic gene sets for metabolic activity analysis and files for Flux balance analysis are available here (https://github.com/wddong1988Mcmaster/STProtocolMouseLiverMetabolism).

## Code availability

All scripts for this pipeline are available on Github: https://github.com/wddong1988Mcmaster/STProtocolMouseLiverMetabolism.

## Acknowledgements

G.R.S. acknowledges the support of a Diabetes Canada Project Grant, a Canadian Institutes of Health Research Foundation Grant, a Tier 1 Canada Research Chair in Metabolic Diseases and a J. Bruce Duncan Endowed Chair in Metabolic Diseases.

## Box 1.

**the minimum requirements for Linux systems**

8-core Intel or AMD processor 64GB RAM

1TB free disk space

64-bit CentOS/RedHat 7.0 or Ubuntu 14.04

The internet is available for machine.

## Box 2.

**Key dataset features and inputs for Space Ranger**

Space Ranger Command Line:

spaceranger count --id=sample \ *#Output directory*

--transcriptome=/ refdata-gex-mm10-2020-A \ *#Path to mouse transcriptome reference*
--probe-set=/Visium_Mouse_Transcriptome_Probe_Set_v1.0_mm10-2020-A.csv \ *#Path to probe set reference*
--fastqs=/fastq_path \ *#Path to FASTQ files from sequencing*
--sample=mysample \ *#Sample name from FASTQ filename*
--cytaimage=/sample_cyt.tiff \ *#Path to CytAssist image from CytAssist equipment (TIFF)*
--image=/sample_scopy.tiff \ *#Path to brightfield image from microscopy (TIFF)*
--slide= Slide_id \ *#Visium slide ID*
--area=A1 \ *#Capture area in Visium slides*
--loupe-alignment=sample.json \ *#Manual image registration file from Loupe Browser*

Notes:

Space Ranger and mouse transcriptome reference are downloaded from https://www.10xgenomics.com/support/software/space-ranger/downloads.

Probe set reference file in CSV format is required since the assay for FFPE tissue samples uses a pair of oligonucleotide probes targeting protein-coding genes. The probe set reference is downloaded from https://www.10xgenomics.com/support/software/space-ranger/downloads.

FASTQ files are generated from sequencing. Sequencing depth for a sample is calculated as following: Coverage percentage of a Visium slide x 5000 total spots x 25,000 read pairs/spot.

A brightfield image is scanned by a microscopy with configuration as following: color camera (3 x 8 bit, 2,424 x 2,424 pixel resolution), white balancing functionality, 2.18 m/pixel minimum capture resolution, exposure times 2-10 milli sec.

An example of Visium slide ID: V43S06-102. This slide serial number is used to retrieve the correct file. An example of capture area: A1 or D1.

JSON files for manual alignment are generated from Loupe Browser as described (https://www.10xgenomics.com/support/software/space-ranger/latest/analysis/inputs/image-cytassist-image-alignment).

Examples for running Space Ranger command line is present in the available code section.

## Box 3.

**Summary metrics result interpretation for liver tissue**

Summary part:

Number of Spots Under Tissue: depends on tissue sample type, for liver >75% coverage is easily achievable.

Mean Reads Under Tissue per Spot: it is recommended >25,000 reads/spot. Median Genes Under Tissue per Spot: it is recommended > 500 genes/spot.

Sequencing:

Number of reads (Total number of read pairs): it is recommended >Coverage percentage of a Visium slide x 5000 total spots x 25,000 reads/spot.

Valid barcodes (Fraction of reads with barcodes that match the whitelist after barcode correction): it is recommended >98.0%.

Valid UMIs (Fraction of reads with valid UMIs): it is recommended >98.0%.

Sequencing saturation (The fraction of reads originating from an already-observed UMI): it is recommended >40.0%.

Q30 Bases in Barcode (Fraction of spot barcode bases with Q-score >= 30): it is recommended >94.0%.

Q30 Bases in Probe Read (Fraction of RNA read bases with Q-score >= 30): it is recommended >94.0%. Q30 Bases in UMI (Fraction of UMI bases with Q-score >= 30): it is recommended >94.5%.

Mapping:

Reads Mapped to Probe Set: it is recommended >96.0%.

Reads Mapped Confidently to Probe Set (Fraction of reads that mapped with MAPQ=255 to one unique probe in the probe set): it is recommended >96.0%.

Reads Mapped Confidently to the Filtered Probe Set: it is recommended >80.0%.

Reads Half-Mapped to Probe Set (Fraction of reads that mapped to unpaired ligation products): it is recommended <1.0%.

Reads Split-Mapped to Probe Set (Fraction of reads that mapped to mispaired ligation products): it is recommended <0.5%.

Spots:

Fraction Reads in Spots Under Tissue: it is recommended >75.0%.

Mean Reads Under Tissue per Spot: it is recommended >25,000 reads/spot. Median UMI Counts Under Tissue per Spot: it is recommended >5,000/spot. Median Genes Under Tissue per Spot: it is recommended >500 genes/spot. Genes Detected (total): it is recommended >19,000 genes.

---

**Extended Data Fig. 1.**
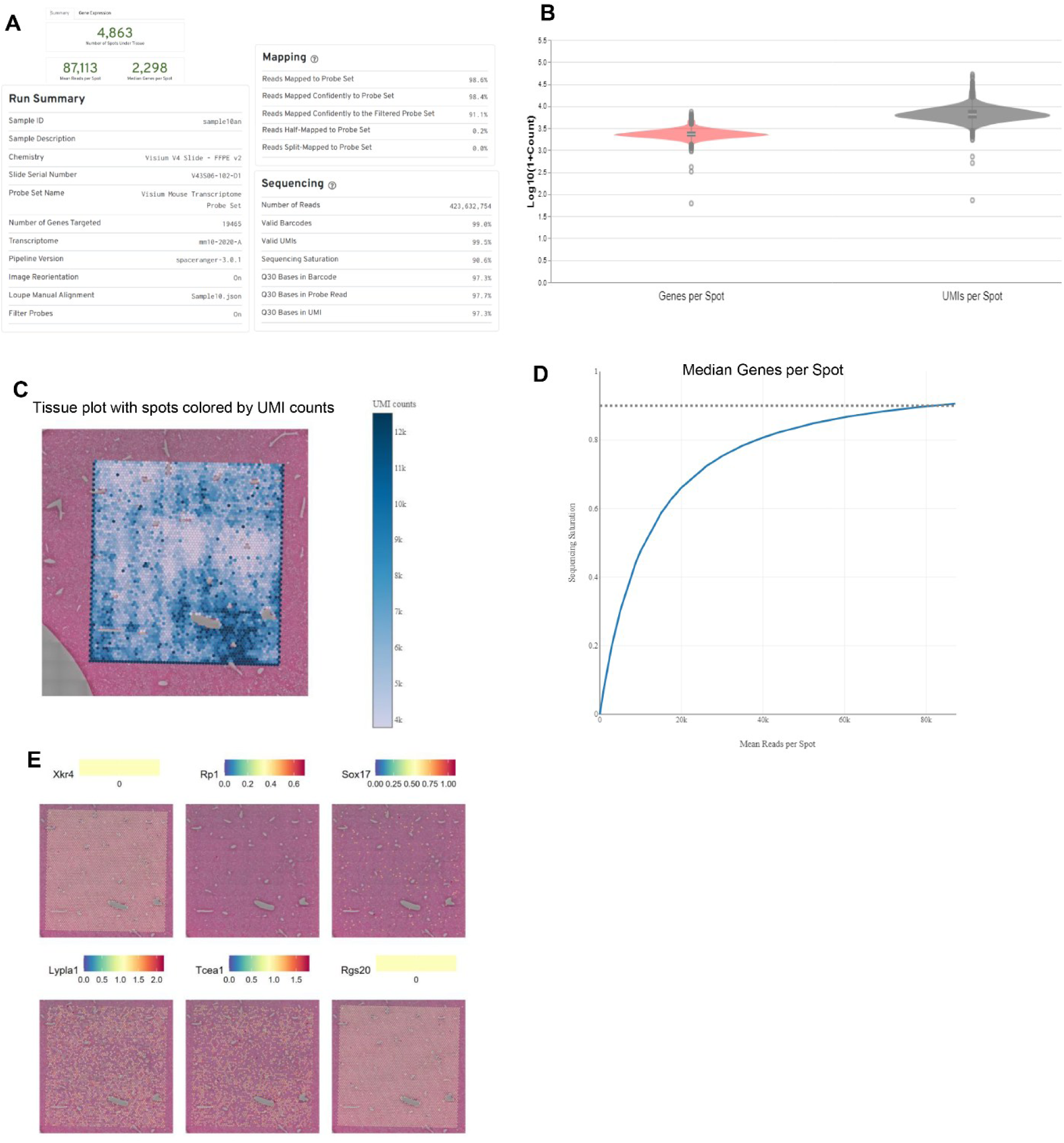
The summary metrics from Space Ranger outputs using to assess sequencing data quality. A, summary parts showing information of mapping and sequencing. B, the number of genes per spot and UMIs per spot. C, heatmap showing the UMI counts per spot on the tissue. D, The median genes per spot. E, the results of spatially variable genes (SVGs) analysis using an annotation-free approach.

