## Supplementary Information for "An Integrated Pipeline for Cell-Type Annotation, Metabolic Profiling, and Spatial Communication Analysis in the Liver using Spatial Transcriptomics"

### Data Generation and Raw data pre-processing

#### *Sample preparation and sequencing*

In this paper, we applied the Visium CytAssist technology. The microarray slides used in Visium contain capture areas composed of numerous spots, each coated with poly-T oligonucleotides, unique molecular identifiers (UMIs), and spatial barcodes[1,2]. These spatial barcodes enable the preservation of transcript localization by capturing mRNA molecules along with their spatial coordinates[1,2]. NGS-based RNA-seq is then used to generate the transcriptomic profiles, with spatial information decoded by mapping the spatial barcodes back to their respective spots on the slide[2]. Two types of microarray slides are available: one featuring 55 × 55 μm barcoded spots separated by 45 μm gaps, and a newer high-resolution version (Visium HD) with 2 × 2 μm barcoded squares arranged without gaps, enabling near single-cell resolution. In this protocol, we focus on the standard Visium slides with 55 μm spatial resolution.

For use with fresh or fresh-frozen samples, tissue is sectioned at 10 μm thickness before being immediately placed onto the capture slide. The tissue is then enzymatically permeabilized to release mRNA, which is captured by the poly-T oligonucleotides on the slide. The optimal permeabilization time must be empirically determined for each tissue type prior to the experiment and imaging options of the tissue samples are limited. For these reasons, FFPE samples are a more common choice due to their availability and compatibility with imaging workflows. For FFPE samples, RNA quality is first assessed, typically requiring a DV200 value > 30%. The tissue is then sectioned at 5 μm thickness, followed by deparaffinization, hematoxylin and eosin (H&E) staining, imaging, and decrosslinking (Fig. 1). This case study includes two experimental conditions (e.g., condition1 vs. condition2), each with biological replicates (condition1a, condition1b et al.). Liver tissues were collected and fixed in 10% neutral-buffered formalin, followed by immersion in 70% ethanol for preservation. The samples were then paraffin-embedded for sectioning.

Additional imaging (e.g., immunofluorescence) can be performed during these steps, provided the reagents are compatible with fixed tissues, thus enabling visualization of protein markers alongside spatial gene expression. Selected tissue sections are transferred onto the capture slide using the Visium CytAssist instrument. Species-specific poly-A-tagged probes are added to the tissues to target the whole transcriptome (up to ~20,000 genes). These probes hybridize to complementary mRNA molecules and are subsequently ligated to confirm hybridization occurred. The ligated probes are then released from the tissue and captured by poly-T probes on the slide, thereby preserving spatial information through attachment of a spatial barcode.

Captured sequences are either reverse transcribed or extended via probe-based methods, then are eluted from the capture oligonucleotides. The resulting material is pooled and amplified then used to generate sequencing libraries which are sequenced at a depth of >25,000 reads/spot under the tissue. Spatial barcodes, incorporated during reverse transcription or probe extension, are used to map sequencing reads back to their original locations on the capture slide. These spatial coordinates are then aligned with tissue images to enable spatially resolved gene expression profiling.

#### *Raw data pre-processing by the Space Ranger pipeline*

To successfully perform raw data processing and quality control, researchers should be comfortable in a Linux-based environment, familiar with command-line tools, and equipped with a computing platform that meets the minimum system requirements outlined in Box 1.

Following sequencing, the first step in the analysis is gene expression quantification (Fig. 1). This process includes manual fiducial alignment of images (optional), read trimming, alignment of transcript reads to the reference genome, barcode correction, and probe filtering (specific to FFPE tissues) to eliminate off-target activity. UMIs are counted to quantify transcript abundance and map the transcriptome to spatial coordinates, resulting in a gene-by-spot expression matrix. Raw data pre-processing is dependent on the underlying spatial transcriptomics platform, as it requires detailed knowledge of the capture slide design. In this protocol, we employ Space Ranger, a dedicated pipeline from 10x Genomics, to perform raw data pre-processing[3]. In this section, we demonstrate how to run the Space Ranger count pipeline on sequencing data derived from an FFPE mouse liver section.

Preparation of materials for running Space Ranger count pipeline (Box 2): To begin, the Space Ranger software package, mouse transcriptome reference files, and probe set reference were downloaded and unpacked. Required inputs for running the Space Ranger count pipeline include: FASTQ files generated from sequencing, a brightfield microscopy image (TIFF), a CytAssist image, the Visium slide ID and its capture area (e.g., A1 or D1), and a JSON file exported from Loupe Browser for manual image alignment. Manual alignment is used to achieve more accurate mapping between the microscopy image and the capture slide.

In our case study, FASTQ files: Generated via sequencing on the Illumina NextSeq 2000 platform using a P3 flow cell with a 2 × 50 bp configuration and a throughput of 1100 million reads.

Brightfield images: Acquired using a Nikon Eclipse 90i upright microscope and saved in TIFF format.

Visium slide ID: V43S05-003 (capture area: A1: condition1a, D1: condition1b), V43S06-102 (A1: condition2a, D1: condition2b).

JSON files: Generated from the Loupe Browser for manual image alignment, following instructions provided by 10x Genomics: https://www.10xgenomics.com/support/software/space-ranger/latest/analysis/inputs/image-cytassist-image-alignment.

The example data files (FASTQ files, brightfield images, JSON files) are available here: https://www.dropbox.com/scl/fo/occj7x04tbkjrnqfrko1b/AOV196ag11OrAIUtWagY6g8?rlkey=qyzhi6f20x517znqovu4n8q22&st=j4mhkfpm&dl=0

Upon successful completion, Space Ranger outputs result in the outs/ subdirectory, including: a run summary metrics (web_summary.html), a Loupe Browser visualization and analysis file (cloupe.cloupe), A matrix of transcript counts in HDF5 format (filtered_feature_bc_matrix.h5), a spatial folder (spatial/) containing high-resolution tissue images and spatial metadata, cluster and differentially expressed genes (DEG) and so on. Among them, the HDF5 matrix file and the spatial/ folder are essential for downstream analysis. The summary metrics (web_summary.html file) are used to assess sequencing data quality. Guidelines for interpreting these quality metrics are provided in Box 3. The file 'web_summary.html' for the one sample is available here (https://github.com/wddong1988Mcmaster/STProtocolMouseLiverMetabolism/blob/main/web_summary10.html). For example, in this dataset, 4,863 tissue-covered spots were detected, with an average of 87,113 reads per spot and 2,298 genes (features) identified per spot. The Run Summary section provides general information about the pipeline execution. The Mapping Results indicate high alignment quality, while the Sequencing Results confirm excellent sequencing performance, with sufficient read coverage to support accurate gene expression quantification (Extended Data Fig. 1A-D). Gene-by-spot expression matrix with spatial information will be generated by this process, which is fundamental for downstream analysis.

#### *QC and normalization*

Once raw data quality has been confirmed, two key outputs from the Space Ranger count pipeline are required for downstream analysis: 'filtered_feature_bc_matrix.h5' and 'spatial' folder (Example files are available here: https://www.dropbox.com/scl/fo/occj7x04tbkjrnqfrko1b/AOV196ag11OrAIUtWagY6g8?rlkey=qyzhi6f20x517znqovu4n8q22&st=j4mhkfpm&dl=0). These files are imported into a Seurat object using the Load10X_Spatial function. We assign sample names to 'orig.ident' and 'cell identity classes' for grouping samples in the following analysis.

In ST, image analysis during preprocessing automatically determines which capture spots contain tissue, reducing the need for extensive QC compared to scRNA-seq, where it is unclear which droplets contain cells[4]. However, technically challenging samples may exhibit variable RNA quality, making additional QC filtering necessary. In addition, initial tissue segmentation is typically generous, often including spots with only partial tissue coverage. This can lead to "edge effects," where spots on the tissue margins cluster together in downstream analyses due to reduced RNA content. To perform QC by filtering spots based on the number of unique genes detected, we first visualize the gene expression distribution across spots (Fig. 3A). Based on this distribution, we define filtering thresholds as follows: 1,500 < features per spot < 7,500. Spots falling below or above this range are excluded to remove low-quality or overly high-expression outliers across all samples. However, in the context of metabolic disease, substantial differences in cell density, such as between fibrotic and non-fibrotic regions, can influence RNA abundance and complicate normalization. Mitochondrial proportion, a common QC metric in scRNA-seq associated with cell damage or apoptosis[5], is generally less informative in ST. This is due to inherent tissue slicing artifacts and the presence of multiple cells within each 55 µm capture spot. Therefore, we do not apply mitochondrial content filtering in our QC workflow. Following filtering, we apply sctransform normalization[6,7], which corrects for technical variability. The glmGamPoi package was used to improve speed. This step returns a Seurat object containing a new assay, named SCT (Fig. 3B).

#### *Dimension reduction*

ST data are high-dimensional and often noisy, as each measured gene represents a separate dimension and carries spatial information. Dimensionality reduction is applied to transform high-dimensional into a low-dimensional representation[8,9]. Dimensionality reduction methods include principal component analysis (PCA), non-negative matrix factorization (NMF)[10], SpatialPCA[8], and Spatial Transcriptomics Analysis with topic Modeling to uncover spatial Patterns (STAMP)[9] and among others[11] (Fig. 2). PCA and NMF are generic algorithms that can be applied to any dataset but do not account for spatial context. In contrast, SpatialPCA and STAMP adapt existing dimension reduction methods PCA and topic Modeling to incorporate spatial information. The incorporation of spatial information generally improves the detection of contiguous spatial domains such as cortical layers of the brain, but it typically reduces the sensitivity to spatially dispersed expression patterns such as tissue infiltrating immune cells. PCA is applied on the normalized and scaled data to do dimensional reduction in this protocol.

Normalized and scaled data are subjected to dimension reduction using PCA via RunPCA function. We plot heatmap by DimHeatmap function to determine which PCs capture the major sources of variation (Fig. 3D). This helps identify PCs that contribute significantly to dataset heterogeneity. To further assess the optimal number of PCs to retain, we use the ElbowPlot function to generate an elbow plot, which visualizes the proportion of variance explained by each PC (Fig. 3E). Selecting the optimal number of PCs for downstream analysis can be challenging. Therefore, it is advisable to repeat the analysis using different numbers of PCs (e.g., 10, 20, 30, or more) to assess how the choice of dimensionality influences the results.

***Clustering***

Clustering methods can be broadly categorized into two classes: statistical approaches (e.g., k-means) and graph-based methods. Graph-based approaches construct a graph—typically a K-nearest neighbor (KNN) or shared nearest neighbor (SNN) graph—and apply community detection algorithms such as Louvain, Leiden, or Smart Local Moving (SLM), or employ deep learning-based techniques such as graph convolutional networks (GCNs)[12]. The Leiden algorithm is a refinement of the Louvain method designed to address limitations such as disconnected communities. It improves both the quality and consistency of clustering by optimizing modularity more effectively[13]. Similarly, the SLM algorithm performs community detection using an advanced local moving heuristic, allowing for more nuanced identification of community structure within the graph[14]. In this protocol, the SNN graph is first constructed by calculating the neighborhood overlap—measured using the Jaccard index—between each spot and its *k.param* nearest neighbors[15,16] using FindNeighbors function[15,16]. Modularity optimization algorithms, such as Louvain, Leiden, or SLM, are then applied to iteratively group spots into clusters based on their network connectivity[17] using the FindClusters function. The resolution parameter in FindClusters controls clustering granularity—higher values result in more clusters. A typical range is 0.4 to 1.2; in this study, we set the resolution to 1.

#### *Non-linear dimensional reduction (UMAP/t-SNE)*

After clustering, the next step is to visualize the data in a lower-dimensional space using non-linear dimensionality reduction techniques (Fig. 2). Unlike the previously used linear method PCA used earlier, which primarily reduces noise and selects principal components for downstream analysis, this step is intended to facilitate data exploration by placing spots with similar gene expression profiles close together in two-dimensional space. Common visualization methods include t-SNE[18] and UMAP[19]. t-SNE models the similarity between data points using a Gaussian distribution in high-dimensional space and maps them into a low-dimensional representation[20]. However, it poorly preserves global structure, meaning the relative positions of clusters may be arbitrary and not biologically meaningful[20]. In contrast, UMAP is able to capture not only fine-grained relationships within clusters but also broader relationships between clusters[19], which may reflect potential physical interactions, shared gene functions, or pathway-level similarities[21]. In this paper, the identified clusters are visualized using UMAP and t-SNE, implemented via the RunUMAP and RunTSNE functions, respectively. The results of UMAP and t-SNE are displayed as two-dimensional scatter plots using the DimPlot function (Fig. 3F and G). In addition, clusters can be visualized on the original tissue image using the SpatialDimPlot function (Fig. 3H). It is essential to interpret the results within the context of known tissue biology, rather than relying solely on computational output.

#### *Identification of cluster biomarkers*

To identify biomarkers for each cluster, DEG analysis is performed using the FindAllMarkers function, which detects genes that are significantly upregulated (positive markers) in a given cluster compared to all other clusters. DoHeatmap function is used to display the top 10 markers for each cluster (Fig. 4A). VlnPlot function is used to visualize expression patterns of selected marker genes in violin plots (Fig. 4B). These markers can also be spatially visualized on the UMAP embedding with the FeaturePlot function (Fig. 4C), or directly on the tissue image using the SpatialFeaturePlot function (Fig. 4D).

#### *Cell type annotation*

Cell type annotation is a fundamental for ST data analysis. Cell identity is not defined by a single gene but by the combinatorial expression of multiple genes at varying levels. Therefore, robust annotation typically relies on a panel of marker genes rather than a single gene. Annotation in ST datasets is particularly difficult due to the limitations of spot-based technologies, such as Visium, which capture RNA from spatial locations without respect to individual cell boundaries[22,23]. To address these challenges, a variety of computational approaches have been developed, including: correlation-based and supervised methods (such as SingleR[24], Scmap[25]), deconvolution-based methods (SpatialDWLS[26], RCTD[27], Cell2location[28]), integrated mapping approaches (Seurat[29]), Spatial domain-based methods (Spatial-ID[23]) with using spatial information, generative pretrained transformers (GPT) based models for automated annotation (GPTCelltype[30]), and novel cell type discovery and annotation tools (STELLAR[31], scBOL[32]). In this case study, a three-step workflow for cell type annotation are applied: automatic annotation using GPT4[30], deconvolution with a scRNA-seq reference, and manual cell type annotation[33].

#### *a. Automatic annotation using GPT*

Automatic cell type annotation is performed as an initial step using GPT4. We utilize the GPTCelltype package to automate cell type annotation by inputting cluster-specific marker gene profiles obtained from Seurat's FindAllMarkers function[30] along with a basic prompt strategy[30] (Fig. 2). Prompt messages are generated based on cluster biomarkers and used to query GPT-4 via the OpenAI API. This process is repeated five times, and the most frequently predicted cell type is selected as the final annotation[30]. The annotation results are visualized on the UMAP using the DimPlot function (Fig. 4E) and mapped onto the original tissue image with the SpatialDimPlot function (Fig. 4F). Although GPT-based annotation is not yet highly accurate, we believe that generative, unsupervised learning models, capable of performing annotations independently of only one/several scRNA-seq reference datasets, represent a significant future direction in the field.

#### *b. Annotation via deconvolution using a scRNA-seq reference*

For sequencing-based ST platforms (e.g., *Visium*), low resolution spots often encompass multiple cells, potentially containing a mixture of different cell types. Even in high-resolution platforms where spot size may be smaller than a single cell, achieving true cellular resolution remains challenging, as mRNA captured in each spot may originate from overlapping cells. Consequently, the transcriptional profile of a single spot often reflects a composite expression pattern from multiple cells, which may obscure true cell-specific gene expression. Cell type deconvolution is commonly employed to predict cell types[29,34]. Deconvolution allows reconstruction of a fine-grained cellular landscape, offering insights into tissue architecture and cellular heterogeneity at near single-cell resolution. We firstly generated a high-quality scRNA-seq reference[35-38] for cell type deconvolution (https://www.dropbox.com/scl/fo/occj7x04tbkjrnqfrko1b/AOV196ag11OrAIUtWagY6g8?rlkey=qyzhi6f20x517znqovu4n8q22&st=j4mhkfpm&dl=0), as visualized on a UMAP plot using the DimPlot function (Fig. 4G). Several methods have been developed for this purpose, such as SpatialDWLS[26], robust cell type decomposition (RCTD)[27], conditional autoregressive-based deconvolution (CARD)[39], anchor-based integration approaches[29]. SpatialDWLS methods are used for deconvolution to identify cell types in spots[29]. SpatialDWLS is able to estimate cell types from ST data via a unique filtering step that removes irrelevant cell types, thereby improving specificity[26]. We then performed normalization, calculated cell type markers, constructed a signature matrix, and executed SpatialDWLS deconvolution within the Giotto framework[40] within the Seurat framework[27]. In this particular case, deconvolution with SpatialDWLS did not substantially improve cell type annotation, but it provided an additional layer of validation. To further refine cell type assignment, we applied RCTD within the Seurat framework. RCTD fits a supervised learning approach to estimate mixtures of cell types at each pixel[27]. We generated an RCTD reference object using the previously constructed scRNA-seq reference and created an RCTD query object from the Seurat spatial transcriptomics data. Deconvolution was then performed using RCTD by creation of the RCTD query object and RCTD deconvolution, and the results were visualized as a bar plot (Fig. 4H). RCTD improved the identification of small cell populations; however, it partially misclassified stellate cells as hepatocytes.

#### *c. Manual annotation*

In this case study, we perform manual annotation at the cluster level using multiple marker genes from FindAllMarker function. Multiple marker genes are used to manually annotate cell types by using different databases such as CellMarker 2.0[41], Tabula Muris[42], PanglaoDB[43], and Datasets from MSigDB[44]. Additionally, scRNA-seq atlases are used as reference points, preferably derived from the same species, organ, and disease context. In this protocol, a custom marker reference was generated for mouse liver with MASLD (https://github.com/wddong1988Mcmaster/STProtocolMouseLiverMetabolism/blob/main/GMT_file/mouse_liver_celltype_markers_MASLD.gmt)[35-38]. Biomarker expression was spatially visualized directly on the tissue using the SpatialFeaturePlot function and further illustrated with violin plots generated using the VlnPlot function. For example, as shown in Fig. 4D, the pericentral hepatocyte markers *Oat* and *Cyp2e1* are highly expressed in spots surrounding the hepatic central veins, indicating that clusters 4 and 5 correspond to pericentral hepatocytes (Fig. 4B). In contrast, the periportal hepatocyte markers *Cyp2f2, Hal* and *Gls2* are predominantly expressed in regions near the hepatic portal vein (Fig. 4I), suggesting that clusters 2 and 8 represent periportal hepatocytes (Fig. 4J).

#### *DEG analysis, pathway enrichment analysis and SVGs*

A common analysis in transcriptomics is the identification of DEGs either between clusters or within the same clusters under different conditions. This approach is used to infer cluster identity or to evaluate gene expression changes in a given cell type across conditions. Methods for DEG analysis include Student's t-test, the Wilcoxon rank-sum test, the limma implementation of the Wilcoxon test, the likelihood-ratio test[45], DESeq2, and spatial linear models incorporating exponential correlation structures[46], among others. In this protocol, we perform DEG analysis using FindMarkers function based on the Wilcoxon rank-sum test. A volcano plot was used to visualize DEGs (Fig. 5A). GO enrichment analysis is conducted to find enriched pathways (Fig. 5B) based on these DEGs.

In this context, SVGs refer to genes whose expression patterns exhibit spatial structure. To detect genes with spatially restricted expression, we perform SVG analysis. Two different approaches were used to perform SVGs analysis. The first approach involves performing DEG analysis based on pre-annotated anatomical regions. For example, we selected regions surrounding the central vein (CV) and portal vein (PV) (Fig. 5C) and performed DEG analysis between these regions (Fig. 5D). An alternative approach is to detect SVG in an unsupervised manner, without anatomical labels, by calculating spatial expression patterns across the tissue. Several computational methods have been developed, including SpatialDE, which uses a Gaussian process framework[47], Splotch[48], SPADE[49] and Trendsceek[50]. In this protocol, we utilize markvario function from the spatstat package, implemented through Seurat as FindSpatiallyVariableFeatures, to identify SVGs. This method is based on marked point processes and detects genes with statistically significant spatial expression patterns[50].

#### *Integrative analysis across multiple samples or conditions*

Integrative analysis aims to remove batch or condition-specific effects while preserving true biological variation. Integration enables the evaluation of cell-type-specific responses to different stimuli, the identification of shared or condition-specific subpopulations, and the discovery of conserved marker genes. Moreover, integrative analysis enhances statistical power and improves the signal-to-noise ratio by pooling information across datasets, effectively increasing the average transcript counts per spot for shared populations. Several integration methods have been developed. These include linear embedding models such as canonical correlation analysis (CCA)[51] and Harmony[52], as well as deep-learning approaches such as scVI[53] (Fig. 2). Harmony projects cells into a shared low-dimensional space, where they are grouped by cell type rather than batch or condition[51,52]. scVI algorithm utilizes a hierarchical Bayesian framework with deep neural networks to model conditional distributions for normalization and analysis[53].

In this section, we demonstrate how to perform integrative analysis across multiple samples or conditions to remove batch- or condition-specific effects while preserving true biological variation. As an example, we use two experimental conditions, each with two biological replicates. The goal of integration is to align the datasets in a shared space to identify common and condition-specific cell subpopulations, and to compare gene expression differences across conditions within each cell type. To achieve this, we first merge the sctransform-normalized datasets using the Merge function. Differential expression analysis is then performed on the merged data using PrepSCTFindMarkers, which utilizes corrected values from individual Seurat objects. Variable features for the merged dataset are generated by combining the variable features identified in each object. Linear dimensionality reduction is then performed using the RunPCA function. Integration is carried out using the IntegrateLayers function, applying either CCA or Harmony. After integration, the standard workflow, including clustering, annotation, and UMAP visualization, is performed as described above. The integration results are visualized using DimPlot (Fig. 5E). Prior to integration, the two samples appear as distinct clusters; after integration, the samples are well-aligned and overlap substantially, indicating effective correction of batch or condition-specific effects and successful integration of shared biological signals. Differential gene expression (DEG) analysis in hepatic stellate cells (HSCs) under two conditions, along with GO enrichment analysis, was performed (Fig. 5F and 5G).

#### *Pseudobulk analysis*

Spots from the same sample typically share a common genetic background and microenvironment, making them pseudoreplicates rather than statistically independent[46,54]. In the pseudobulk analysis, raw counts from all spots belonging to the same cluster within a sample are aggregated using the AggregateExpression function, resulting in one expression profile per cell type per sample. Additionally, generalized linear mixed models with random effects for samples can be used to further address the conservative nature and reduced sensitivity associated with pseudobulk aggregation[54]. DEG analysis is then conducted using the FindMarkers function with the DESeq2 method as shown in a volcano plot (Fig. 5H). GO enrichment analysis is further performed to detect functional differences in a dot plot (Fig. 5I).

#### *Cellular communication*

A variety of computational tools have been developed to study cell-cell communication by integrating ligand-receptor interactions with spatial proximity. These include CellChat[55], CellPhoneDB (with built-in ligand–receptor databases)[56], node-centric expression models (NCEM) based on graph neural networks[57], spatial variance component analysis (SVCA) using Gaussian processes[58], SpatialDM[59] and Tensor-cell2cell using tensor decomposition[60]. In this protocol, we apply CellChat, a tool that leverages network analysis and patter recognition techniques, to analyze spatially resolved cell-cell communication[55]. Using the CellChat tool enables identification of novel communication networks between distinct cell types and signaling pathways within the tissue microenvironment. To investigate cellular communication in liver tissues, we performed cellular communication analysis across multiple spatial transcriptomics datasets using CellChat, which integrates gene expression data with spatial proximity. After loading the data, a CellChat object was created for each sample. Using a curated ligand–receptor interaction database, CellChat predicts intercellular communication events. The cell-communication analysis includes the number and strength of interactions (Fig. 5L- 5N), visualization of the network using a circle plot and a spatial image overlay (Fig. 5O, P), network centrality scores (Fig. 5Q), and spatially visualized in tissue images (Fig. 5R), contributions of individual ligand–receptor pairs (Fig. 5S), the expression levels and spatial distributions of representative ligands (e.g., *Cd44*) and receptors (e.g., *Cd74*) (Fig. 5T).

#### *Metabolic activity analysis*

#### *a. Metabolic pathway activity*

Metabolic activity can be evaluated using GSEA[61] with predefined gene sets representing specific metabolic pathways. We generated a curated collection of mouse-specific metabolic gene sets. The collection includes metabolic pathways derived from KEGG, as well as other metabolic pathways in the liver, such as de novo lipogenesis (DNL), very low-density lipoprotein (VLDL) export, fatty acid oxidation (FAO), cholesterol synthesis, liver inflammation and fibrosis (GMT file: https://github.com/wddong1988Mcmaster/STProtocolMouseLiverMetabolism/tree/main/GMT_file). To improve the reliability of pathway activity inference, we applied integrated gene set enrichment analysis (irGSEA), which aggregates results from multiple scoring methods (AUCell, UCell, singscore, ssGSEA, and JASMINE), using the robust rank aggregation (RRA) algorithm[62]. This integrative approach enhances the stability and robustness of metabolic pathway activity scoring. Differences in metabolic activity patterns across liver cell types, pathway–gene relationships and the activity of specific pathways were evaluated based on GSEA.

#### *b. Metabolic interactions*

Metabolic interactions are critical for coordinating cellular function, because cells exchange nutrients, compete for metabolic substrates, and secrete metabolites that influence the function of neighboring cells. These interactions reshape cellular behavior, modulate the tissue microenvironment, and define immune niches. Metabolic crosstalk can be investigated through correlation analysis of metabolic pathway activity scores across cell types. Another approach involves inferring intercellular communication via ligand–receptor gene pairs associated with metabolic processes. In this protocol, we also curated ligand-receptor interactions associated with glucose, lipid, and amino acid metabolism from the CellChat database. We then used these curated pairs to perform cell–cell communication analysis focused specifically on metabolic signaling. We first measured metabolic interactions by analyzing correlations between pathway activities. ssGSEA[63] was performed using our curated metabolic gene sets to calculate enrichment scores. Spearman correlation analysis was then used to evaluate relationships between metabolic pathways across different cell types and experimental conditions. The correlations between glycolysis/gluconeogenesis and TCA cycle in hepatocytes and pathway correlations under different conditions were measured (Fig. 6F and Fig. 6G).

As an alternative approach, we examined cellular communication mediated by metabolic signaling. To do this, we curated ligand–receptor signaling pathways associated with glucose, lipid, and amino acid metabolism from the CellChat database[55]. Pathways related to glucose metabolism include INSULIN, IGF, GIPR, GCG, LEP, ADIPONECTIN, RESISTIN, and ApoE. Those involved in lipid metabolism include LEP, ADIPONECTIN, RESISTIN, ANGPTL, ApoE, ApoA, ApoB, LXA4, 27HC, cholesterol, calcitriol, desmosterol, DHEA, DHT, estradiol, progesterone, and testosterone. Pathways related to amino acid metabolism include GABA-A, GABA-B, glutamate, glycine, serotonin-dopamine, histamine, IGFBP, NMU, NPY, NTS, VIP, PACAP, somatostatin, TAFA, and PROK. To evaluate these pathways, we quantified interactions, analyzed a specific signaling pathways, such IGF signaling (Fig. 6H-6K). The most influential cell types and pathways were identified by network centrality analysis (Fig. 6L). We also demonstrated how to compare intercellular interactions between conditions (e.g., control vs. treatment or healthy vs. diseased) by assessing the number and strength of interactions, metrics analysis, communication networks of multiple ligand–receptor pairs (Fig. 6M-6O).

#### *c. Flux balance analysis (FBA)*

Several tools have been developed to perform FBA, including Compass[64] and METAFlux[65]. GEMs are typically structured around two key components: a stoichiometric matrix and gene-protein-reaction (GPR) rules, which link genes to enzymatic activities. Commonly used GEMs include Recon[66] and Human1 for human metabolism, and iMM1415[67] and Mouse1[68] for mouse models. Among these, Recon2 provides a more comprehensive reconstruction of human metabolism than currently available mouse models. In this case study, we adapted Recon2 to perform FBA in mouse data, leveraging the high degree of conservation in core metabolic pathways between humans and mice. To demonstrate how to assess metabolic programs using Compass[64], we exported the gene expression matrix and metadata after downsampling to 300 cells per cell type. To perform FBA, we utilized the high-quality genome-scale metabolic model (GEM)--RECON2, which encompasses central metabolic pathways that are highly conserved between humans and mice. FBA was conducted on the ST data using Compass[64] to generate penalty scores for each reaction in each cell, producing output in reaction TSV files. To compare the interaction difference between conditions and cell types, Wilcoxon rank-sum test was applied (Fig. 6P-6R).
